# Decreased hippocampal neurite density in late middle-aged adults following prenatal exposure to higher levels of maternal inflammation

**DOI:** 10.1101/2024.10.01.616156

**Authors:** Raana A. Mohyee, Blake L. Elliott, Madeline R. Pike, Emma Smith, Ann M. Kring, Ingrid R. Olson, Elizabeth C. Breen, Barbara A. Cohn, Piera M. Cirillo, Nickilou Y. Krigbaum, Thomas M. Olino, Mark D’Esposito, Ashby B. Cogan, Bhakti P. Patwardan, Lauren M. Ellman

## Abstract

In animal models, exposure to heightened maternal inflammation in utero is associated with altered offspring hippocampal development, including reduced dendritic arborization and density. However, the effects of prenatal maternal inflammation (PNMI) on offspring hippocampal microstructure in humans remains unclear. Here, we examined the relationship between exposure to PNMI and neurite density in the hippocampus and its subfields among offspring during late middle age. Participants included 72 mother-offspring dyads from the Child Health and Development Studies (CHDS) cohort. Data for four inflammatory biomarkers (IL-6, IL-8, IL-1 receptor antagonist [IL-1RA], and soluble TNF receptor-II [sTNF-RII]) were available from first and second trimester maternal sera. Neurite density in the offspring hippocampus and its subfields was estimated using microstructural modeling of offsprings’ diffusion-weighted Magnetic Resonance Imaging data (mean age of offspring at imaging = 59 years; 51% male). We estimated the relationship between each biomarker and region-of-interest’s neurite density. Higher first trimester maternal IL-1RA and IL-6 levels were associated with lower offspring hippocampal neurite density. These relationships were specific to the CA3, CA4, dentate gyrus, and subiculum subfields. In addition, higher second trimester IL-6 was associated with lower subiculum neurite density. Our findings reveal that exposure to heightened prenatal levels of maternal inflammation is linked to altered offspring hippocampal microstructure in late middle age, which could have implications for memory decreases during this period and may be relevant for understanding risk of aging-related cognitive changes.

**Significance Statement:** The contribution of prenatal maternal inflammation (PNMI) to offspring brain microstructure in later life is well established in animal models but poorly understood in humans. Our study discovered long-lasting impacts of elevated PNMI during early mid-gestation on the structural integrity of the hippocampus in offspring during late middle age. Our findings underscore the potential role of prenatal insults in aging-related neurological and cognitive decline, as the observed degradation in hippocampal microstructure is present over half a century following exposure.

## Introduction

During fetal development, a cascade of biological events results in the formation of distinct cortical and subcortical brain structures, which have been shown to be influenced by maternal health and the prenatal environment (1–4). A growing body of literature implicates the role of prenatal maternal immune functioning, especially systemic inflammation during pregnancy, in several neurological and behavioral outcomes in offspring throughout development (5). Studies have associated prenatal maternal inflammation (PNMI) with increased prevalence of autism spectrum disorder (5–9), psychotic disorders (10–12), depression (13), and childhood internalization and externalization symptoms (14, 15), disorders and sequelae that have a demonstrated origin in altered neurodevelopment (5).

Importantly, alterations in the structure and function of the hippocampus, a subcortical region integral to learning and memory, emotion, and regulation of physiological stress responses (16–22), have been implicated in the etiology and course of many of these psychiatric conditions, as well as types of dementia including Alzheimer’s disease (23–25). The hippocampus, which completes much of its cytoarchitectural maturation and subfield differentiation during the second and third trimesters of pregnancy, is particularly sensitive to neuroimmune activity during the fetal period (26–29). The hippocampus contains some of the highest densities of microglia and several proinflammatory cytokine receptors in the brain, especially the interleukin-1β (IL-1β) receptor (27, 30).

Furthermore, the hippocampus is one of few brain structures that remains highly structurally and functionally plastic throughout adulthood (31). As such, the hippocampus serves as a prime avenue for exploring the long-term developmental outcomes associated with prenatal exposure to heightened maternal inflammation (26, 32, 33).

Studies using animal models for maternal immune activation (MIA) have linked heightened exposure to MIA to several offspring hippocampal alterations, including increases and decreases in volume, changes in dendritic structure and complexity, decreased myelination, and decreases in pre- and post-synaptic proteins (reviewed in 26, 34–36). In preterm infants exposed to Histological chorioamnionitis, PNMI was associated with increased CNS expression of proinflammatory cytokines, including IL-6 and IL-1β (a trending, but not statistically significant relationship for IL-1β) and decreased myelination (36). Another study of neonates identified similar relationships between TNF-α and measures of diffusivity in several major white matter tracts in the brain (37).

Further, the structural changes observed following PNMI differ across the subfields of the developing hippocampus, each of which has unique histological and functional characteristics (38). Animal studies have shown that PNMI decreases neuron density and the number of pyramidal cells in the Cornu Ammonis (CA) subregions 1, 2, and 3 (39, 40), reduces CA1 axonal size and myelin thickness (41), and impairs cell proliferation and neurogenesis in the Dentate Gyrus, leading to memory deficits (42–44). Furthermore, PNMI results in reduced CA3 fiber density (45), aberrant CA3-to-CA1 synaptic transmission (46), and altered excitatory postsynaptic potentials in CA1 pyramidal neurons, resulting in abnormal object information processing (47). These findings highlight the need to investigate whether PNMI differentially affects specific hippocampal subfields to better understand its impact on neuronal functioning and behavior.

In addition, both clinical and preclinical studies demonstrate the importance of timing of PNMI exposure in relation to offspring neurodevelopment, though findings differ somewhat across species. Studies on rodents and non-human primates have found that alterations in offspring brain development after PNMI largely depend on the timing of exposure (26). For example, a study of rhesus monkeys demonstrates that early gestational MIA resulted in increases in brain volume, whereas MIA introduced later in gestation resulted in decreases in brain volume (reviewed in 26). Several studies have also implicated the importance of timing of PNMI exposure in the development of offspring behavioral phenotypes. For example, one rodent study has linked late gestational exposure to PNMI to behavioral deficits mirroring negative symptoms of schizophrenia, as well as altered dopaminergic, GABAergic, and glutamatergic neurotransmission in the hippocampus (48). Another rodent study found that key cellular processes affecting neuronal architecture, including neurogenesis, reelin immunoreactivity, and apoptosis, were more drastically altered in offspring following PNMI during middle gestation than in late gestation, with various behavioral deficits observed following exposure at both timepoints (49). Several studies in rodents have shown that several behavioral abnormalities analogous to those observed in human psychiatric disorders, including ultrasonic vocalizations, asociality, social recognition deficits, repetitive or stereotypic behaviors, and memory deficits, are all affected by timing of exposure to MIA (34). However, a study in non-human primates that compared first- and second-trimester exposure to MIA, did not find any significant differences between first- and second-trimester-exposed groups in similar behavioral assays (35). In PNMI-exposed infants of adolescent mothers, both IL-6 and C-reactive protein, another marker of immune activity, were correlated with diffusion magnetic resonance imaging (MRI) changes in a number of subcortical regions and parts of the medial temporal lobe (50). In the present study, we will investigate timing effects by examining PNMI during both the first and second trimesters.

Along with timing of PNMI exposure, preclinical studies suggest that offspring age at the time of outcome measurement provides another source of variation in existing preclinical findings on PNMI and hippocampal development, although this has yet to be explored in human research. For instance, PNMI-related changes in offspring expression of several immunogenic chemosignals, both peripheral and intracranial, including in the hippocampus specifically, have been observed in proximal stages of development, but surprisingly not in adulthood (34, 51). Furthermore, PNMI has been shown to impair hippocampal neurogenesis in juvenile rodents; yet, changes in neurogenesis observed in adulthood, as a result of PNMI, appear to be spatially restricted and sex-specific (33, 52, 53). One recently published epidemiological study demonstrated that the effects of PNMI on human hippocampal and memory functioning observed in middle life emerge as a consequence of reproductive aging in sex-specific patterns (54). Another study examining the brains of PNMI-exposed human offspring across childhood and adolescence observed that many of the differences in brain structure observed early in childhood may resolve themselves through “catch-up growth” during this highly dynamic period of brain development, suggesting the need for follow-up examination of these neurodevelopmental outcomes across the lifespan (55). Determining the association between PNMI and offspring hippocampal microstructure in later stages of human development is crucial given the relevance for aging-related memory decline and neurodegenerative diseases in this age group (56).

Very few human studies have investigated the effects of PNMI on hippocampus microarchitecture, or indeed other brain structures, past the neonatal period, giving rise to questions regarding the translation of preclinical findings across the human lifespan (49, 57). Moreover, existing human studies have either depended on indirect measures of PNMI (58), utilized postmortem methods (59), or used nonspecific metrics of structural change, like total volume or diffusion tensor-derived metrics (60, 61). However, diffusion tensor derived metrics assumes that water diffusion has a Gaussian distribution, but biological tissue displays non-Gaussian diffusion due to the restricted diffusion by barriers such as organelles and cell membranes (62). Additionally, diffusion tensor derived metrics fail to accurately describe microstructural changes in gray matter because water diffusion is relatively isotropic (63). To overcome these challenges, we collected multishell diffusion magnetic resonance imaging (MRI) and implemented a biophysical tissue model, the Neurite Orientation Dispersion and Density Imaging (NODDI). The NODDI model uses diffusion MRI to measure density and spatial distribution of the orientation of neurites (e.g., dendrites, axons, and their branches) (64). The NODDI model was developed to overcome challenges in measuring the microstructure of heterogeneous tissues in the brain such as gray matter structures, lending itself to the study of the hippocampus particularly well.

To our knowledge, no human studies have investigated the effects of PNMI on the structure of the hippocampus in late middle age in a population-based sample. Consequently, in a sample of late middle age human adults who have been followed since the fetal period as part of a longitudinal cohort (Figure 1), we investigated the association between PNMI on adult offspring’s hippocampal microstructure using noninvasive, *in vivo* brain imaging and novel microstructural modeling techniques (i.e., NODDI). We also sought to investigate PNMI’s effects on cytoarchitecture in the hippocampal subfields. We hypothesized that heightened PNMI would be associated with reduced neurite density in the hippocampus of adult offspring. Additionally, we hypothesized that these effects would be specific to the CA1, CA3, and dentate gyrus.

**Figure 1.**
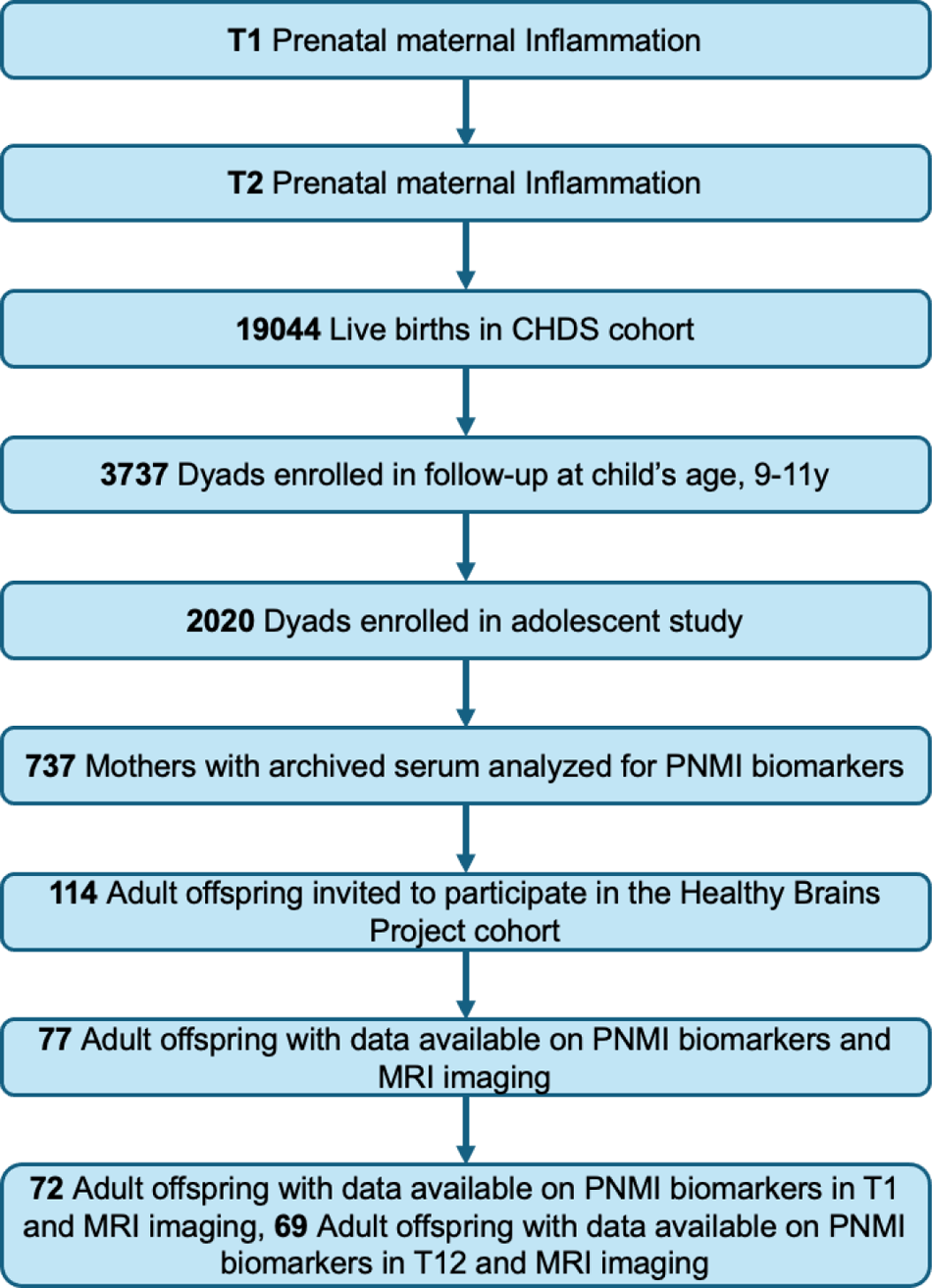
Participants Retained in Subsamples of the Child Health and Development Studies (CHDS) and Healthy Brain Project (HBP). T1 = first trimester of pregnancy; T2 = second trimester of pregnancy.

## Results

### Descriptives and Bivariate Correlations with Covariates

Demographic information is presented in Tables 1 and 2. Violin plots showing the distributions of ICVF in the hippocampus and its subfields in this sample and violin plots for each trimester, grouped by cytokine can be found in Figures 2 and 3. Bivariate correlations for first and second trimester variables are reported in Supporting Information Tables S1 and S2. We first conducted bivariate analyses using Spearman’s correlations between our covariates of interest, PNMI biomarkers, and hippocampal data. Covariates (maternal education, offspring intracranial volume, offspring age at time of imaging, and offspring sex) were included which have been implicated in the focal processes (i.e. hippocampal neurite density or PNMI) (16,58,59). Maternal education and intracranial volume were not significantly correlated with ICVF, nor with any prenatal biomarkers at trimester 1 [T1; mean (SD) gestation = 11.21 (2.50) weeks] nor trimester 2 [T2; mean (SD) gestation = 23.17 (2.37) weeks]. Offspring sex (female) was negatively correlated with CA1 and CA3 ICVF. Offspring.

**Figure 2.**
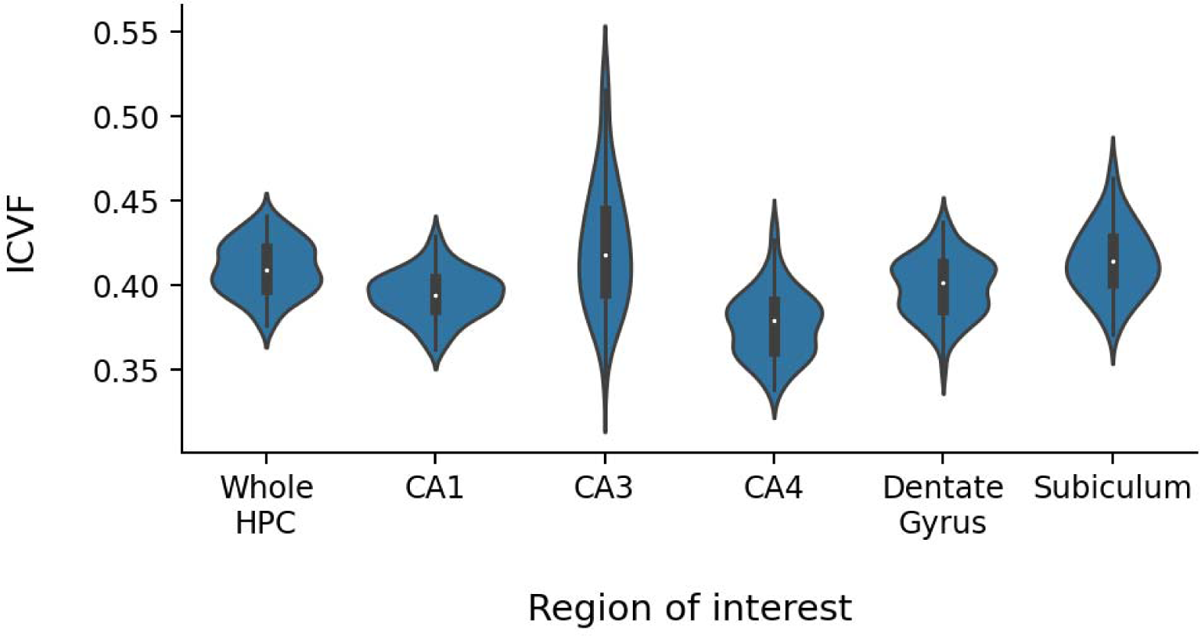
Neurite density (ICVF) across the hippocampus and subfields. The y-axis represents neurite density measured using intracellular volume fraction (ICVF). The x-axis denotes different hippocampal subfields, including the whole hippocampus (Whole HPC), Cornu Ammonis 1 (CA1), Cornu Ammonis 3 (CA3), Cornu Ammonis 4 (CA4), the dentate gyrus (DG), and the subiculum (SUB). Each violin plot displays the distribution of ICVF values within the respective region, where the white dot represents the median, the thick black bar indicates the interquartile range (IQR, 25th–75th percentile), and the thin black whiskers extend to 1.5 times the IQR range.

**Figure 3.**
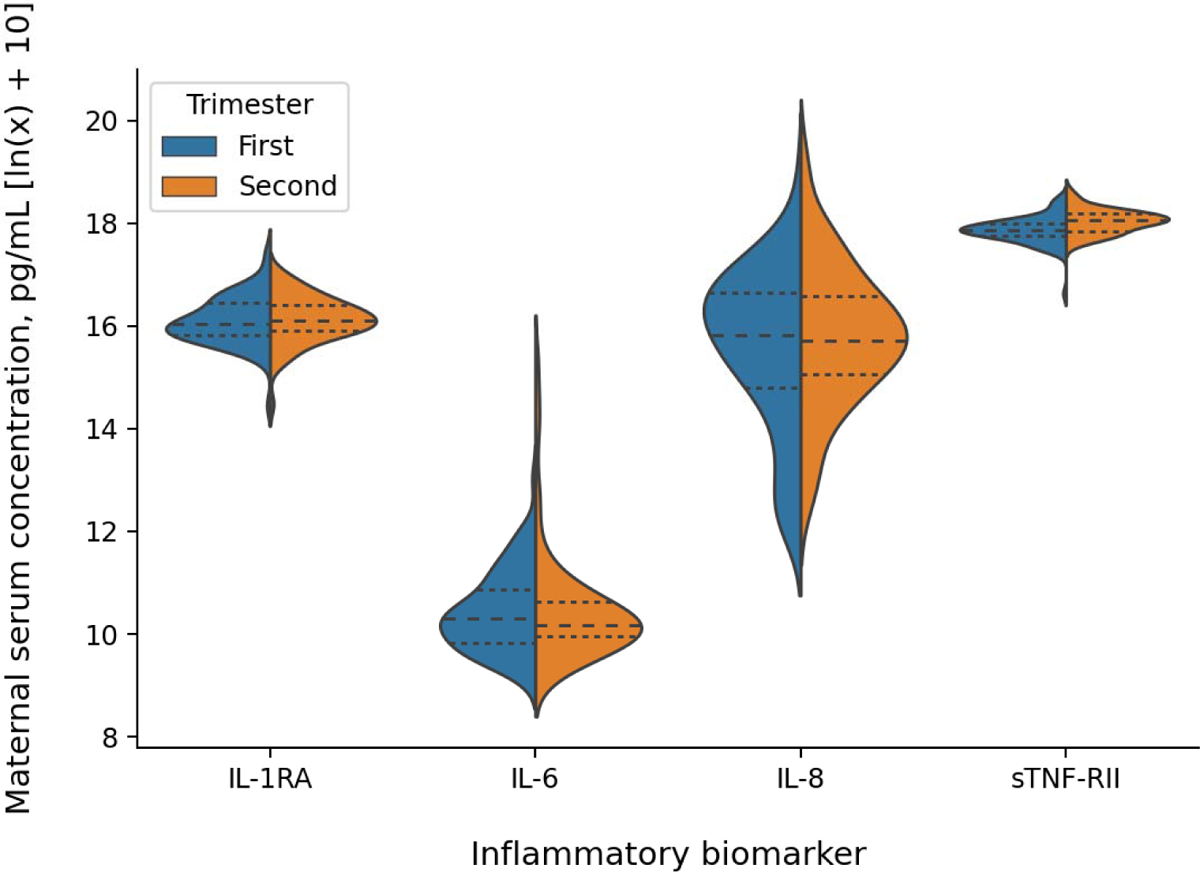
Violin plots of maternal serum levels of inflammatory biomarkers for each trimester. First-trimester values are shown on the left (blue), and second-trimester values are shown on the right (orange) for each biomarker. The y-axis represents maternal serum concentrations (pg/mL), which have been transformed using ln(x) + 10. The central dashed line within each violin plot represents the median, while the dotted lines indicate the interquartile range (25th–75th percentiles). The violin shapes illustrate the kernel density estimation of the data distribution for each trimester.

**Table 1.** Participant Demographics.

|  |  |
| --- | --- |
| **Age at interview** (years) M (SD) [range] | 59.6 (1.1) [57.3-62.3] |
| **Sex** % (n) Male | 48.6 (36) |
| **Ethnicity** % (n) Hispanic | 8.1 (6) |
| **Race** % (n) |  |
| American Indian/Alaska Native | 2.7 (2) |
| Asian | 6.8 (5) |
| Black | 12.2 (9) |
| White | 70.3 (52) |
| More than one race | 6.8 (5) |
| Unknown | 1.4 (1) |
| **Gestational age at birth** (weeks) M (SD) [range] | 40 (1.72) [33.1-44.1] |
| **Birthweight** (grams) M (SD) [range] | 3354.5 (516.6) [1956.2-4677.8] |

*Note.* N = 74; demographics are from all participants who were included in at least one hierarchical regression analysis; T1 = trimester one; T2 = trimester two.

**Table 2.** Maternal Demographics.

|  |  |
| --- | --- |
| **Age at offspring birth** (years) M (SD) [range] | 29.2 (6.4) [18-43] |
| **Race** % (n) |  |
| □ White | 74.3 (55) |
| □ Black | 13.5 (10) |
| □ Asian | 6.8 (5) |
| □ Mexican/Hispanic | 5.4 (4) |
| **Education** % (n) |  |
| < high school | 6.8 (5) |
| = high school | 33.8 (25) |
| > high school | 59.5 (44) |
| **Pre-pregnancy BMI** (kg/m<sup>2</sup>) M (SD) [range] | 21.7 (3.3) [16.7-37.9] |
| **Parity** M (SD) [range] | 2.1 (1.9) [0.0-8.0] |
| **Serum T1 IL-6** (pg/mL) M (SD) [range] | 2.2 (2.7) [0.5-20.6] |
| **Serum T1 IL-8** (pg/mL) M (SD) [range] | 611.5 (911.1) [8.0-6807.3] |
| **Serum T1 IL-1ra** (pg/mL) M (SD) [range] | 502.6 (269.5) [85.9-1775.4] |
| **Serum T1 sTNF-RII** (pg/mL) M (SD) [range] | 2705 (698) [759-4839] |
| **Serum T2 IL-6** (pg/mL) M (SD) [range] | 6.6 (26.2) [0.5-197.6] |
| **Serum T2 IL-8** (pg/mL) M (SD) [range] | 809.5 (1544.9) [12.2-10323] |
| **Serum T2 IL-1ra** (pg/mL) M (SD) [range] | 505.1 (217.3) [88.7-1137.6] |
| **Serum T2 sTNF-RII** (pg/mL) M (SD) [range] | 3181 (744) [1884-5735] |

*Note.* N = 74; demographics are from all mothers of participants who were included in at least one hierarchical regression analysis. Serum cytokine levels are reported in pg/mL for ease of interpretation but were natural log transformed for all analyses.

Offspring sex (female) was negatively correlated with CA1 and CA3 ICVF. Coincidentally, offspring age at the time of imaging was negatively correlated with maternal serum IL-8 at T1, though this relationship does not lend itself to meaningful interpretation.

### Prenatal Maternal Inflammation and Hippocampal Neurite Density

We next conducted analyses to determine whether individual differences in hippocampal ICVF were related to maternal inflammatory biomarkers using hierarchical multiple regression. Parameter estimates can be found in Table 3. At T1, higher maternal IL-1β, as measured by serum concentrations of its receptor antagonist (IL-1RA, see Methods for more information) and IL-6 were associated with lower offspring ICVF in the hippocampus. At T2, we found no significant associations between any biomarker and the whole hippocampus.

**Table 3.** Results of hierarchical linear regression analyses for prenatal maternal inflammation and offspring hippocampal microstructure during late middle age.

| Independent Variable | | $\beta$ | SE | $t$ | $p$ | $\Delta R^2$ |
| --- | --- | --- | --- | --- | --- | --- |
| **Step 1: Covariates** |  |  |  |  |  | 0.040 |
| Sex |  | -0.181 | 0.004 | -1.438 | 0.155 |  |
| Age |  | -0.084 | 0.002 | -0.827 | 0.411 |  |
| Intracranial Volume |  | -0.163 | 1.210e-08 | -1.590 | 0.117 |  |
| Maternal Education, = High School |  | 0.522 | 0.008 | 2.160 | **0.034\*** |  |
| Maternal Education, > High School |  | 0.458 | 0.007 | 1.976 | 0.052 |  |
| **Step 2: Additive Effect** |  |  |  |  |  |  |
| First Trimester | IL-1ra | -0.269 | 0.004 | -2.333 | **0.023\*** | **0.061\*** |
|  | IL-6 | -0.281 | 0.002 | -2.444 | **0.017\*** | **0.067\*** |
|  | IL-8 | -0.087 | 0.001 | -0.672 | 0.504 | 0.029 |
|  | sTNF-RII | -0.091 | 0.006 | -0.762 | 0.449 | -0.006 |
| Second Trimester | IL-1ra | -0.122 | 0.004 | -0.990 | 0.326 | -0.001 |
|  | IL-6 | 0.088 | 0.002 | 0.691 | 0.492 | -0.008 |
|  | IL-8 | 0.012 | 0.001 | 0.094 | 0.925 | -0.015 |
|  | sTNF-RII | 0.107 | 0.009 | 0.852 | 0.398 | -0.004 |

*Note.* Each of the additive effects were assessed using separate models; IL = Interleukin, RA = receptor antagonist, sTNF-RII = soluble tumor necrosis factor receptor-II, HPC = hippocampus, CA = Cornu Ammonis, DG = Dentate Gyrus (including the molecular and granule cell layers), \* $p < 0.05$ .

We next investigated whether the relationship between PNMI and hippocampal ICVF was specific to particular hippocampal subfields (depicted in Figure 4B) using hierarchical multiple regression. Parameter estimates can be found in Table 4. When individual hippocampal subfields were analyzed, at T1 both IL-1RA (Figure 4C) and IL-6 (Figure 4E) were associated with lower offspring ICVF in the CA3 and dentate gyrus subfields (Figure 5). Additionally, higher IL-1RA showed an uncorrected association with lower subiculum subfield ICVF and higher IL-6 showed an uncorrected association with lower CA4 subfield ICVF, although these did not survive multiple comparison correction. There were no significant correlations between any biomarker and the CA1 subfield. At T2, we found that IL-6 showed an uncorrected association with lower offspring ICVF in the subiculum, but this did not survive multiple comparison correction. No other associations were observed between any biomarker and any hippocampal subfield at T2 (Figure 4D, F).

**Figure 4.**
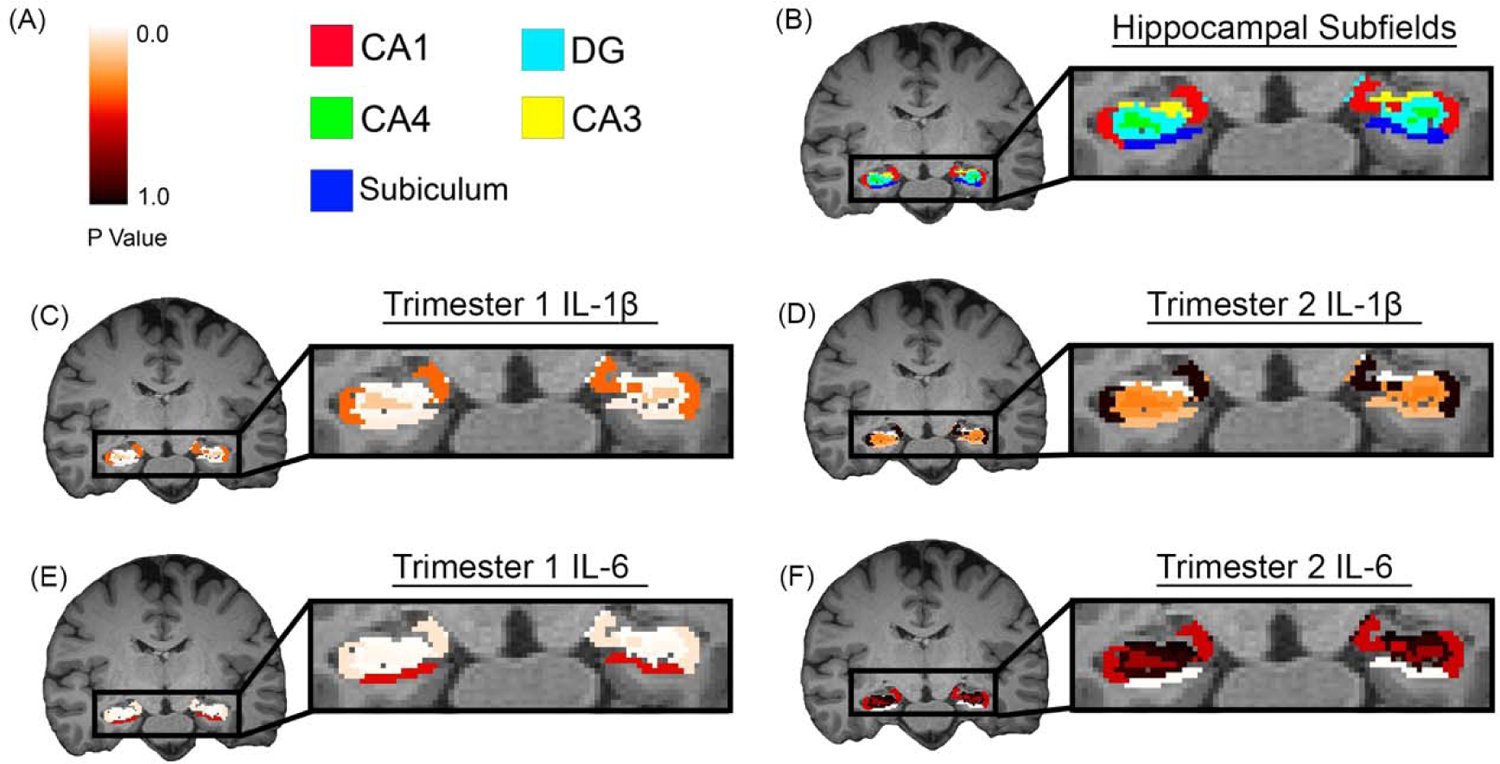
Associations between hippocampal subfield neurite density (ICVF) and prenatal immune biomarkers. (A) Color maps for B-F; (B) map of subfields of the hippocampus; (C) significance levels of relationship between prenatal exposure to first-trimester maternal IL-1β and subfield neurite density in the first trimester; (D) significance levels of relationship between prenatal exposure to second-trimester maternal IL-1β and subfield neurite density; (E) significance levels of relationship between prenatal exposure to first-trimester maternal IL-6 and subfield neurite density in the first trimester; (F) significance levels of relationship between prenatal exposure to second-trimester maternal IL-6 and subfield neurite density in the second trimester; CA = Cornu Ammonis, DG = Dentate Gyrus, Molecular Layer, and Granule Cell Layer.

**Figure 5.**
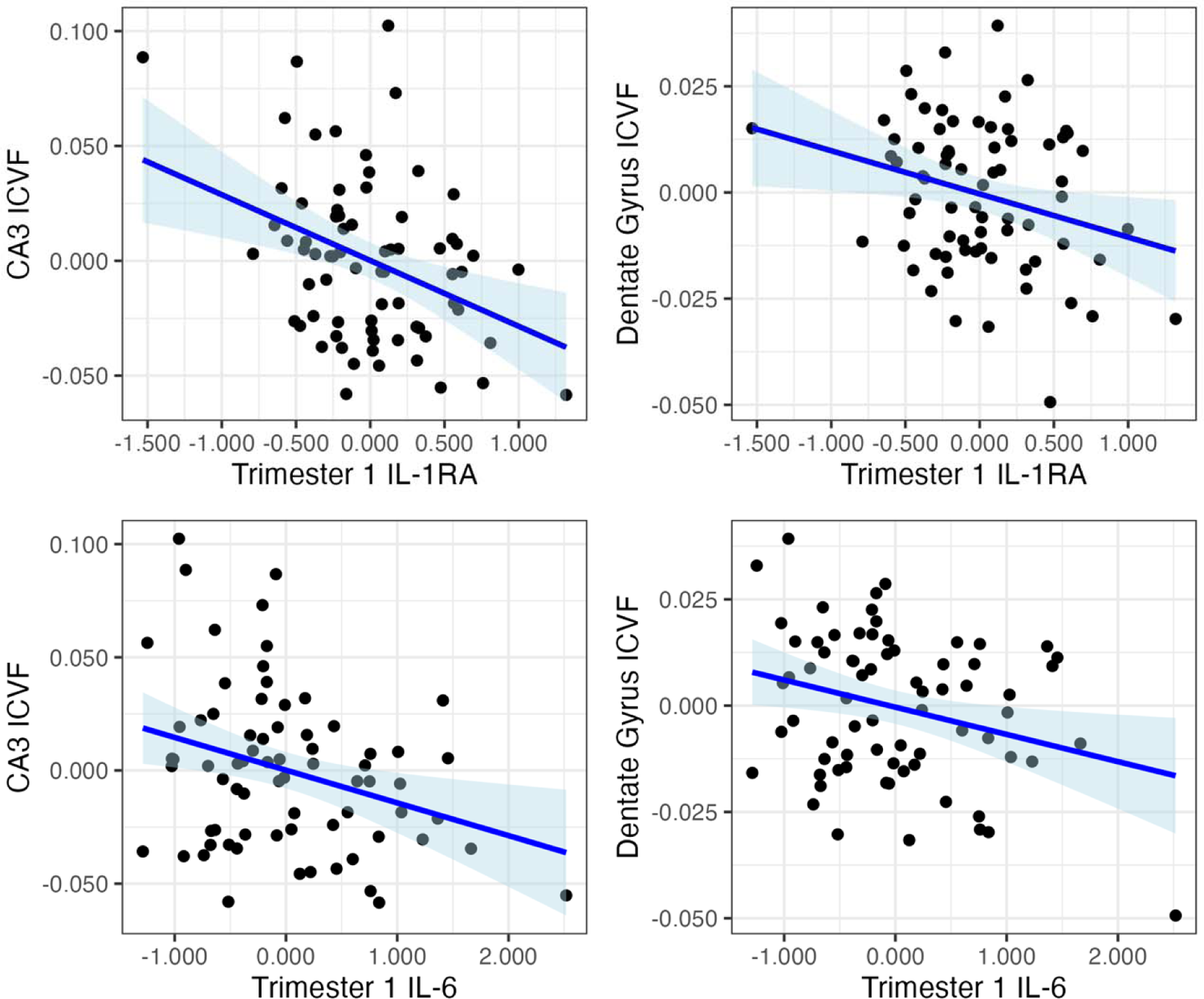
Relationship between maternal serum inflammatory cytokines in early pregnancy and hippocampal subfield neurite density (ICVF) in middle age. The x-axis represents residualized inflammatory cytokine levels (natural-log-transformed pg/mL) after adjusting for intracranial volume, age, maternal education, and sex. The y-axis represents residualized hippocampal subfield ICVF values, controlling for the same covariates. Each scatter plot shows individual data points (participants), with the solid blue line indicating the linear regression fit and the light blue shaded area representing the 95% confidence interval.

**Table 4.**
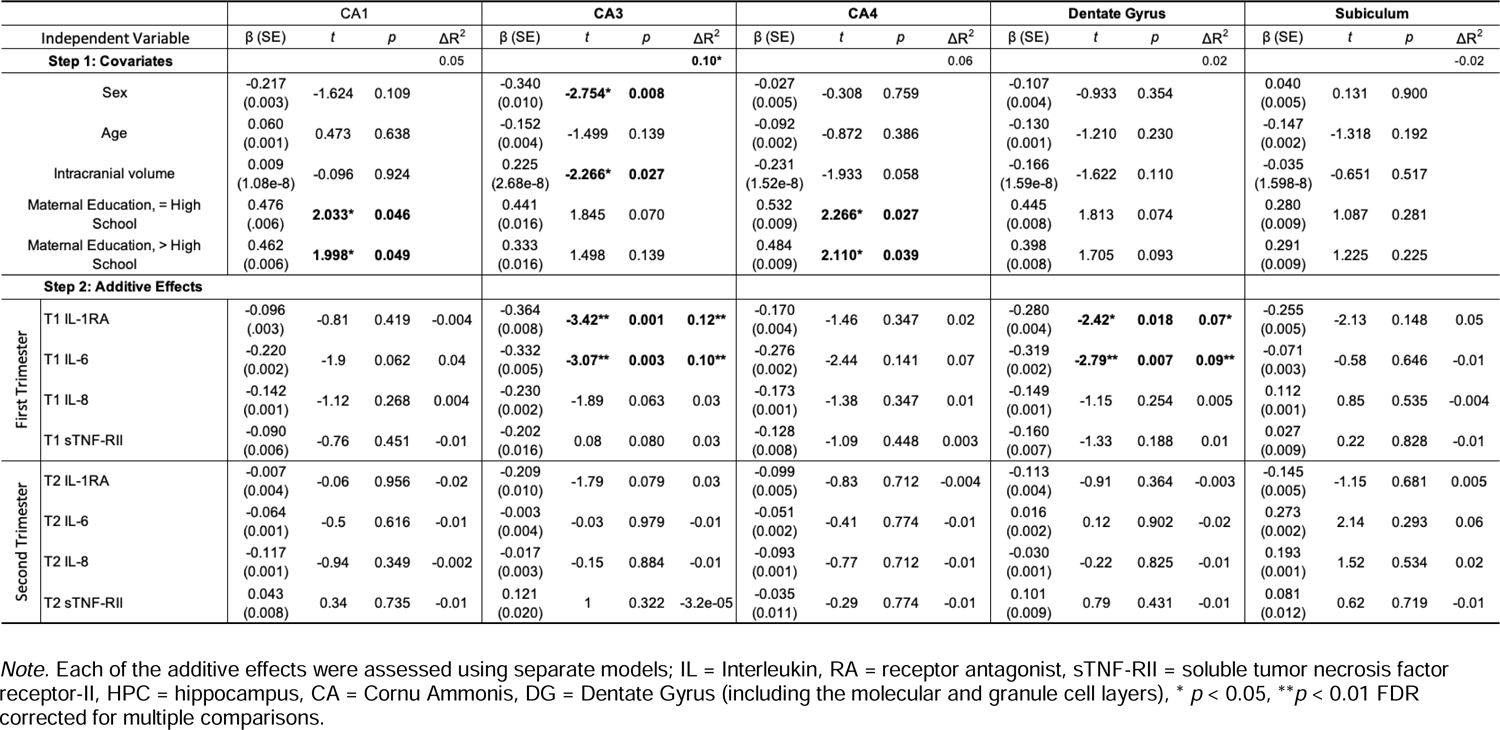
Results of hierarchical linear regression analyses for prenatal maternal inflammation and offspring hippocampal microstructure during late middle age for each hippocampal subfield.

## Discussion

To our knowledge, this is the first human study to find an association between exposure to heightened prenatal inflammation and offspring neurite density in the hippocampus and its subfields during late middle age. We found that higher levels of maternal inflammatory biomarkers, specifically IL-1RA and IL-6 during the first trimester, were associated with reduced ICVF in the hippocampus and certain hippocampal subfields, including CA3, CA4, dentate gyrus, and subiculum (although the CA4 and subiculum subfields did not survive multiple comparison correction). Interestingly, no significant relationship was observed in the CA1 subfield, and only IL-6 from the second trimester showed an uncorrected association with the subiculum subfield. These results indicate that PNMI is associated with reduced neurite density in the hippocampus and that these effects depend on the timing of exposure, are distributed differentially across subfields of the hippocampus, and are evident in late middle age. These findings are consistent with previous studies that demonstrate microstructural alterations in the hippocampus of nonhuman animals following PNMI (33, 53, 65).

Our results implicate heightened prenatal exposure to IL-1β (via measurement of IL-1RA), in altered development of the offspring hippocampus. Research has shown that IL-1RA is a strong, valid, and reliable estimate of IL-1β activity that is present at much higher concentrations in sera than the cytokine itself in pregnancy, which is necessary for improving detectable values of cytokine activity in the prenatal period (14, 15, 66). Extensive evidence from animal models of maternal prenatal immune activation have demonstrated reduced hippocampal neurogenesis in maternal IL-1β-exposed offspring (33). An *in vitro* study of human hippocampal cells identified upregulation of kynurenine production and gene expression for several related neurotoxic enzymes following experimental exposure to heightened IL-1β, demonstrating at least one mechanism of action through which the observed effects on hippocampal neurite density may have been borne out by heightened exposure to prenatal, maternal IL-1β (67). Furthermore, IL-1β is an integral molecular mediator of the cellular processes in the hippocampus that support long-term memory consolidation, the expression of which in the healthy hippocampus is tightly regulated (25, 68). Importantly, blocking IL-1β signaling in models of Alzheimer’s Disease reduces tau pathology, microglial activation, and secretion of cytokines by astrocytes (68).

Additionally, in vivo human studies have linked IL-1β to cognitive abilities. In a prospective birth cohort in the Southeastern United States, heightened maternal IL-1β during pregnancy was negatively associated with several cognitive abilities including overall intelligence and nonverbal and spatial abilities [measured with the Differential Abilities Scale [DAS-II]; (69)], in offspring in early childhood (70). The hippocampus may play a crucial role in these abilities.

Traditionally the hippocampus has been shown to be critical in the formation of episodic memories, spatial memories, and the ability to navigate through space (16, 71). However, the hippocampus has now been shown to be crucial for many broader aspects of cognition, such as perception, working memory, semantic search, concept formation, future planning, and social behavior (72–80). While IQ and broader cognitive abilities such as nonverbal reasoning engage multiple brain regions, including the hippocampus, our results suggest that the observed associations may be driven, at least partially, by the hippocampal contribution to these specific domains. Together, these findings suggest that neurite density is sensitive to cytokine-specific effects on human hippocampal cytoarchitecture and that these microstructural changes observed in the hippocampus in response to heightened IL-1β expression could drive functional impairments related to learning, memory, and other cognitive domains at both early and later stages of development.

The significant relationship between neurite density in offspring hippocampi and maternal IL-6 from both trimesters (though evidence in the second trimester was only associated with the subiculum subfield prior to multiple comparisons corrections) is consistent with the extensive preclinical findings implicating IL-6 in hippocampal development and function. Similar to IL-1β, IL-6 has been associated with the maintenance of long-term potentiation, a cellular process key to the consolidation of long-term memory (27), and appears to be a key molecular mediator of symptoms of neuropsychiatric disorders in which hippocampal abnormalities have been regularly observed (81, 82). Elevated maternal prenatal IL-6 in rodents has been shown to upregulate glutamate receptor expression, associated with a significant increase in excitatory postsynaptic currents, in the hippocampus of offspring, causing an imbalance of excitatory and inhibitory activity in the region (83). This imbalance likely has downstream effects on the cytoarchitecture of the hippocampus. Exposure to heightened maternal IL-6 in utero has been linked to several subfield-specific microstructural changes in offspring hippocampi, including decreased cell proliferation and neuronal survival in the granule cell layer of the dentate gyrus, neuronal loss in the hilus region of the dentate gyrus, and decreased neuronal density in the CA1 subfield of female offspring and in CA2 and CA3 subfields of the male offspring (33). As such, our findings with IL-6 confirm not only the importance of the cytokine in the developing hippocampus but also highlight the potential regional specificity of subfields of the hippocampus most heavily involved in learning and memory function, namely the dentate gyrus and CA3 subfields.

The dentate gyrus and CA3 subfields, both of which have been shown to be particularly susceptible to microstructural changes following PNMI in the present study, are intricately anatomically connected and are jointly critical for memory function. The dentate gyrus receives dense input from the entorhinal cortex via the perforant pathway and connects with the CA3 subfield via the mossy fiber pathway. The CA3 subfield has reciprocal connections back to the dentate gyrus, which are critical for memory function (84). The dentate gyrus is thought to play a central role in pattern separation, a process that distinguishes and decorrelates incoming input patterns from the entorhinal cortex, ensuring the encoding of distinct memories. The CA3 region is believed to contribute to pattern completion, aiding in the retrieval of memories by associating and reinstating stored information (85–91). APOE ε4, the best-known genetic risk factor for Alzheimer’s Disease, has been shown to have a selective effect on the dentate gyrus and CA3 subfields (92, 93). Importantly, our sample ranged from 57 to 63 years old, which is when the first signs of cognitive decline start to emerge (94). Indeed, recent research has demonstrated that heightened PNMI was associated with altered hippocampal functioning in the brain during a verbal memory task, hindered task performance, and increased immune activity in middle age (54). Thus, this study’s findings indicating a strong association between heightened PNMI and reduced neurite density in the DG and CA3 subfields offer valuable insights into the potential long-term consequences of maternal prenatal inflammation on cognitive health in aging and neurodegenerative disease risk, emphasizing the need for further translational research into the underlying mechanisms driving these specific associations.

Additionally, our findings may provide some insight into recent studies linking PNMI to offspring depression and related cognitive impairments. Previous research has demonstrated that higher concentrations of prenatal maternal IL-1RA and IL-6 are associated with higher adolescent depressive symptoms (15), and the effect of IL-1RA on depressive symptoms was shown to be partly mediated by childhood cognitive performance (95). As depression is stress-sensitive and the hippocampus is highly responsive to stress (24, 96–98), it may serve as a key region connecting PNMI and offspring depression.

Further, chronic inflammation, especially low-grade or systemic inflammation, has been increasingly implicated in the development and progression of cognitive decline, including conditions like Alzheimer’s disease and other forms of dementia (99, 100). IL-6 has been particularly highlighted, with high levels of systemic IL-6 associated with higher rates of cognitive decline (99). Animal studies have provided foundational evidence that abnormal glial– neuron interaction in the hippocampus may contribute to the pathogenesis of cognitive disturbances after neuroinflammation (101). Thus, the current finding of increased prenatal IL-1RA and IL-6 associated with decreased offspring hippocampal neurite density (particularly in CA3 and dentate gyrus regions) could be critical for understanding the neurobiological mechanisms contributing to depression and memory disorders, suggesting that the origins of these relationships could begin before birth.

In this sample, we found no significant associations with hippocampal or subfield neurite density with measures of sTNF-RII, a valid and reliable estimate of TNF-α activity in pregnancy, from either trimester. TNF-α plays an important role in modulating hippocampal-dependent cognitive functions, including sleep and memory function (68). Furthermore, TNF-α demonstrably alters hippocampal microstructure through direct regulation of hippocampal synaptic expression of AMPA and GABA receptors (25, 102), and heightened expression of TNF-α has been shown to reduce cell proliferation and neuronal differentiation in the hippocampus (32). This robust relationship between TNF-α and hippocampal function has largely been characterized by the cross-sectional study of endogenously produced TNF-α in the brain, and current research findings suggest that this cytokine may not be essential to producing the behavioral deficits typically observed in maternal immune activation models (102). However, recently published findings from another prospective birth cohort study show that higher gestational exposure to maternal TNF-α was associated with poorer memory performance and with differences in hippocampal activity during the performance of memory tasks, as measured by functional MRI, in sex-specific patterns in middle-aged offspring (54). Notably, these findings resulted from analyses of the relationship between maternal serological cytokine concentrations drawn from the third trimester of pregnancy and analyzed cytokine levels instead of soluble receptor levels (the latter representing a better estimate of cytokine activity), which may have accounted for the differences in results. As such, more work examining how the timing-, sex-, and maternal TNF-α activity levels influences offspring hippocampal development is needed to draw firmer conclusions about the role of this particular cytokine.

Despite mixed evidence supporting different mechanisms of action for the translation of inflammation from mother to offspring in utero, findings from preclinical studies increasingly converge on the possibility of altered development of offspring microglia as a notable underlying mechanism driving the neuroimmunological changes observed in offspring as a consequence of PNMI (27, 68, 81). Some evidence also supports the idea that peripheral cytokines expressed by the mother can cross or upregulate cytokine production by the placenta, which then enter the fetal bloodstream (32, 43, 103, 104); however, further human studies would be necessary to parse out specific mechanisms of action.

The prominence of first trimester exposures in the significant relationships observed between PNMI and hippocampal microstructure in this study and others points to the potential mediating role of offspring microglia. The first trimester is the period during which microglia populate the burgeoning central nervous system in the embryo, and evidence suggests that exposure of developing microglia to dysregulated cytokine expression results in the accelerated maturation and morphological activation of these cells, such that when exposed to a secondary immune insult, these cells mount stronger and more rapid cellular responses (27, 105). It is possible that PNMI alters the susceptibility of microglia-mediated developmental processes in the hippocampus, like decreased neurogenesis postnatally, resulting in lasting changes in hippocampal microstructure (27, 68, 106). Indeed, Graciarena et al. (44)found that the impaired neurogenesis in the dentate gyrus of adult mice prenatally exposed to maternal inflammation was associated with increased activation of microglia in particular. Thus, the long-term impact of PNMI on hippocampal microstructure observed in this study could be partially related to a cascade of alterations to neurodevelopment, with offspring microglia activation contributing to neurodevelopmental sequelae.

In addition to the potential mediating role of offspring microglia activation, a second alternative explanation for our results may be found in the placenta. Specifically, susceptibility of the placenta to maternal inflammation during pregnancy likely alters prenatal development through a number of physiological pathways independent of the offspring’s immune system, including disruption of the flow of oxygen or nutrients to the fetus (107). Furthermore, maternal inflammation during pregnancy can succeed a number of lifestyle and environmental factors that alter offspring hippocampal development through several non-neurodevelopmental pathways (30). Thus, an alternative interpretation of this data could entertain PNMI as a proxy measure for other physiological and psychological insults experienced by the mother driving the observed changes in offspring hippocampal development through further alternative pathways.

### Strengths, Limitations and Future Directions

This study’s strengths include the use of a unique cohort with prenatal serum data *and* modern neuroimaging data collected during late middle age. The availability of trimester-specific, serologically defined biomarkers of PNMI enables an investigation of its systemic immunological effects. Despite the age of our samples, we successfully detected biomarkers in all cases and focused on analytes present at higher concentrations in pregnant serum. Additionally, our study implemented the innovative Neurite Orientation Dispersion and Density Imaging (NODDI) model with diffusion imaging. As demonstrated by Zhang et al. (2012), NODDI provides greater specificity than fractional anisotropy (FA) in diffusion imaging (64). Voxels with distinct NODDI parameters can produce identical FA values, making them indistinguishable using FA alone. Our use of this innovative technique offers a more detailed understanding of hippocampal structure in late middle age, providing insights into the intricate neural mechanisms affected by PNMI.

Though the current study discovered later-life impacts of elevated maternal prenatal inflammation on hippocampal cytoarchitecture, this study has several limitations to consider. The quantification of maternal inflammation for a study like this poses several challenges. Given the complex immunological changes experienced during each stage of pregnancy (108), the single measurements of maternal cytokine expression from each trimester offer limited insight into maternal immune function in this sample. Additionally, degradation of the samples is always possible; however, multiple steps were taken to ensure that degradation of samples did not influence cytokine analyses (see 14 for more details) and there is no reason to believe that degradation of samples would occur in a directional capacity to influence results. Furthermore, it was beyond the scope of this study to investigate potential sources of maternal inflammation, such as infection (109), stress (110), obesity (111), nutrition (112), and/or toxic exposures (113), which could all exert additional independent and/or interactive teratogenic effects on brain development, limiting the conclusions about environmental factors leading to increased PNMI.

Immune biomarkers from offspring were not available in this study, such that little information about how PNMI relates to offspring immune functioning, and how offspring immune functioning, in turn, relates to the microstructural changes observed, could be gleaned from this sample. Future studies would benefit from finer-grain data capturing the developmental trajectories of both the hippocampus and immune system of offspring together. Furthermore, there could be widespread effects of PNMI throughout the cortex and subcortically beyond the hippocampus. Although an interesting avenue for future research, the absence of offspring immune data does not negate our findings. Given the temporal precedence of prenatal inflammation, any downstream immune phenotype in the offspring would most plausibly serve as a moderator or mediator of the observed associations, rather than a confounder or an alternative explanation. Additionally, both preclinical and human studies suggest that prenatal inflammation can induce inflammatory states in offspring, and that both prenatal and postnatal inflammation can lead to alterations in the hippocampus (26, 114–116). Importantly, it has also been shown in preclinical models that prenatal maternal inflammation leads to hippocampal deficits in the absence of persistent offspring inflammation across the lifespan (116). Although this is one of the largest neuroimaging studies of a birth cohort of this kind, constraints on study design and sample size still only allowed for modest power to detect significant associations and, therefore, examining interactions and other complex relationships between variables of interest was not possible (117).

Additionally, though these analyses controlled for offspring sex, the design of this study, as well as limited sample size, preclude rigorous examination of sex differences in changes to hippocampal microstructure following PNMI in this sample. Several studies in rodents have characterized differential effects of PNMI across male and female offspring (33, 48, 118). Importantly, a recent study published using data from a birth cohort recruited from the Northeastern United States demonstrated that heightened PNMI was associated with altered memory circuitry functioning in the brain, including the hippocampus, reduced verbal memory performance, and increased immune activity in middle age, in a manner highly dependent on both offspring sex and reproductive age, with postmenopausal women experiencing the worst outcomes (54). Future studies should prioritize quantifying these sex-specific and -dependent developmental differences in relation to aging-related changes in the brain and behavior to inform public health interventions for concerns such as the increasingly asymmetrical burden of Alzheimer’s disease and other dementias, which have been observed more often in women than in men across many studies (119).

Lastly, it is important to acknowledge the limited generalizability of these results. There are very few studies that have examined the long-term neural consequences of prenatal inflammation in human offspring, and only a handful of cohorts worldwide are equipped to address this question. This makes our study and cohort uniquely suited to provide a rare and valuable first step in understanding the long-term sequelae of prenatal inflammation. However, our cohort is derived from a sample from the San Francisco Bay Area, a geographic region uniquely affected by the changing cultural landscape of the mid-to-late twentieth century, during which prenatal data collection was conducted, and which individuals from diverse demographic backgrounds experienced very differently. Moreover, while the CHDS was representative of the Bay Area at the time, it was not representative of the diverse population of both this region and the greater national population currently. Thus, our findings are best interpreted as a mechanistic or etiologic proof-of-concept. Replication in more diverse and contemporary samples is necessary to determine how contextual factors such as socioeconomic status, race/ethnicity, and access to healthcare may moderate the long-term effects of prenatal inflammation.

## Conclusion

Our study represents a significant advancement in our understanding of the enduring effects of PNMI on offspring hippocampal microstructure in later life in humans. The large cohort, combined with the longitudinal nature of our study, with retention rates allowing for analysis decades after pregnancy, have illuminated effects of maternal prenatal inflammation on hippocampal microstructure in a period of late middle age (late 50’s-early 60’s) that represents a key time for early identification and intervention for cognitive decline. By revealing a specific link between heightened maternal prenatal inflammation and alterations in offspring neurite density within key hippocampal subfields, our research highlights the potential implications of programming on aging-related neurological and cognitive decline. These insights underscore the importance of further investigation into the mechanisms underlying these associations and their broader implications for public health.

## Materials and Methods

### Participants

Participants were recruited from the Healthy Brains Project, a follow-up study of the Child Health and Development Studies (CHDS). CHDS is a prospective multi-generational cohort originally recruited for the purpose of studying health outcomes across the life span following prenatal exposure to environmental factors that affect human development (N = 19,044 live births) (120). This cohort enrolled pregnant women who were living in the greater Alameda County, California, United States area, between 1959 and 1966, and who received prenatal care from medical facilities within the Kaiser Foundation Health Plan.

Adult offspring who participated in the CHDS follow-up during adolescence (see 14 for details of this study), and for whom archived maternal serum samples from T1 and/or T2 had been previously analyzed for markers of inflammation (n=737), were invited to participate in the Healthy Brains Project adult follow up study. Recruitment followed a systematic, tiered sampling method and prioritized individuals who resided within close enough proximity to the neuroimaging facility (Berkeley, California) to participate in the neuroimaging portion of the study. Current analyses drew from the 114 individuals enrolled in the Healthy Brains Project cohort at the time of present analyses (circa January 2023). Of these 114 cohort members, 85 had completed the MRI portion of the study. Three participants from whom high-resolution MRI scans of the hippocampus, T1-weighted structural MRI scans, or diffusion-weighted MRI scans were not collected were excluded from analysis. An additional five participants whose MRI scans were deemed too low quality after visual inspection were excluded. From the remaining 77 participants, 5 were missing T1 sera data, and 8 were missing T2 sera data, leaving a final sample of 72 participants with T1 sera data and full MRI and 69 participants with T2 sera data and full MRI. 72 participants with MRI and T1 sera data, and 69 participants with MRI and T2 sera data were included in the final analyses (Figure 1).

The research protocol was approved by the Institutional Review Boards (IRBs) at Temple University, University of California, Berkeley, and the Public Health Institute of California and all adult offspring enrolled in the Healthy Brains Project provided full informed written consent. However, the original CHDS predated IRBs; therefore mothers and offspring in the original CHDS voluntarily participated in in-person interviews and gave oral permission to researchers for medical record access for themselves and their children.

### Demographics Variables

Detailed sociodemographic information was collected for all mothers during the second trimester of pregnancy (mean gestation = 15.3 weeks, SD 6.8 weeks). Maternal education was used as a proxy for postnatal advantages because it 1) was shown to be correlated with other measures of socioeconomic status (SES; e.g., annual household income) in similar published studies, 2) has been previously used in this way by multiple similar studies (121), 3) had the most complete data of the available SES measures in this dataset, and 4) accounts for variations in hippocampal functioning related to maternal education. Maternal education was categorized as “Did not complete High School,” “Completed High School,” or “Completed more than High School’’ and coded as a factor variable for analyses. Offspring race was categorized as “American Indian/Alaska Native”, “Asian”, “Black”, and “White.” Offspring sex assigned at birth (herein referred to as “sex”) was categorized as “Male” or “Female.” See Tables 1 and 2 for more detailed demographic information.

### Prenatal Maternal Inflammation (PNMI)

Prenatal Maternal Inflammation (PNMI) was quantified from archived prenatal first [T1; mean (SD) gestation = 11.21 (2.50) weeks] and second [T2; - mean (SD) gestation = 23.17 (2.37) weeks] trimester maternal serum as previously described in detail (see 14). PNMI biomarker assays were conducted using high-sensitivity (IL-6, IL-8) and regular sensitivity (sTNF-RII, IL-1RA) ELISAs (R&D Systems, Inc., Minneapolis, MN), and raw concentrations were reported in picograms per milliliter (pg/mL).

### MRI Data acquisition

MRI scans were acquired on a 3 Tesla Siemens MAGNETOM Trio scanner using a 32-channel head coil at the University of California, Berkeley, Henry H. Wheeler Jr. Brain Imaging Center. A T1-weighted structural image was acquired using multi-echo, magnetization-prepared, 180-degree radio-frequency pulses and rapid gradient-echo (MEMPRAGE) sampling (TR=2,530.0 ms, TE=1.64-7.22 ms, FOV=256.0 mm, flip angle=7°, slice thickness=1.0 mm, slices per slab=176.0, voxel size=1.0 × 1.0 × 1.0 mm, matrix size=176 × 256 × 256 voxels, scan time [min]=6:03). Additionally, a high-resolution, T2-weighted scan with partial brain coverage focused on the medial temporal lobe and an oblique coronal slice orientation (positioned orthogonally to the longitudinal axis of the hippocampus) was acquired from each participant for use in parcellation of the subfields of the hippocampus (TR= 8,020.0 ms, TE= 51.0 ms, FOV=175 mm, flip angle=122°, slice thickness=1.5 mm, slices=40, voxel size=0.4 × 0.4 × 1.5 mm, matrix size=448 × 448 × 40 voxels, scan time [min]=3:54). Diffusion-weighted images were collected along 111 directions using a multi-shell sequence (b = 1000 s/mm2 and b = 2000 s/mm2) and sensitivity encoding image reconstruction (SENSE) (TR=3,400.0 ms, TE=94.8 ms, FOV=210 mm, flip angle=78°, slice thickness=2.0 mm, slices=69, voxel size=2.0 × 2.0 × 2.0 mm, matrix size=104 × 104 × 69 voxels), scan time [min]=6:31). Additional b0 (no diffusion weighting) images were collected in the reverse phase encoding direction for estimation of susceptibility-induced distortions in image preprocessing (described in subsequent sections).

### MRI Processing

#### Hippocampal Subfield Segmentation

Region-of-interest (ROI) masks were generated for the following subfields of the hippocampus using T1-weighted and high-resolution, T2-weighted structural scans with FreeSurfer Version 7.2.0: CA1, CA3, and CA4; the dentate gyrus; and the subiculum (122). Due to the difficulty in reliably segmenting CA2 and CA3 separately, the Free Surfer algorithm opts to subsume CA2 in the segmentation for CA3, an approach we adhered to in this study. For the purposes of these analyses, the dentate gyrus, granule cell layer, and molecular layer segmentations were combined, as cytoarchitectural delineation of the subfields of the hippocampus suggest that the molecular and granule cell layers are structures within the dentate gyrus (123). The three subregions of the subiculum were similarly analyzed together as a single structure.

#### Diffusion-weighted Imaging (DWI)

Diffusion-weighted scans were denoised using dwidenoise from MRtrix3 Version 3.0.2 (124–127). Susceptibility-induced distortions were then calculated using top-up from FSL Version 6.0.5.1 (128, 129). Eddy current distortion correction and susceptibility-induced distortion correction were performed with slice-to-volume motion correction and outlier replacement using eddy from FSL Version 6.0.5.1(130–132). The distortion-corrected volumes were then visually inspected for remaining image quality issues before NODDI metrics were estimated.

#### NODDI Metrics

The NODDI model was fitted to participants’ preprocessed diffusion images using the AMICO pipeline Version 1.4.2 to produce estimates of neurite density for the whole brain (133). The NODDI model hybridizes the tensor-based and ball-and-stick modeling approaches to consider three contributions to the overall diffusion signal: intra-neurite fluid, modeled as ‘sticks’, or highly-restricted diffusion; fluid in cell bodies and the immediate extracellular space, modeled as anisotropic, or partially restricted diffusion; and free water, modeled as isotropic diffusion (64). As previously stated, NODDI calculates the relative contributions of each component of the tissue to produce a whole-brain map of intra-neurite fluid, termed the Intracellular Volume Fraction (heretofore abbreviated as ICVF). The ICVF has been validated against gold-standard histological measures of neurite density. ICVF has shown a positive correlation with myelin expression (134), and dendritic complexity and length (135), and highly negative correlation with measures of cell body density (134). Therefore, ICVF has been interpreted as the density of neurites (axons and dendrites), based on intracellular diffusion, and is also referred to as the neurite density index (59, 136). The average ICVF was computed bilaterally for each hippocampal subfield for use in the statistical analyses.

#### Data Analysis Plan

A priori power analyses were conducted to detect small-to-moderate effect sizes (f² =.15), which indicated a required sample size of 55. Our final sample of 72 participants, therefore, provide sufficient power to detect small to medium effects—effect sizes that are typical in neuroimaging studies (137, 138).

All analyses were conducted in R version 4.2.1 using a combination of base R and tidyverse functions (139, 140). The dependent variables were examined for normality by examining skewness and kurtosis values, and visually, through inspection of graphic representations of these data. All independent variables (PNMI biomarkers) were natural log-transformed due to skewed values (as is typical with these biomarkers) (14, 15, 95).

Bivariate analyses using Spearman’s correlations were conducted between the biomarkers of PNMI and hippocampal data (141). Covariates (maternal education, offspring intracranial volume, offspring age at time of imaging, and offspring sex) were identified as either variables significantly associated with both the independent and dependent variables or variables implicated in the focal processes (i.e., hippocampal neurite density or PNMI).

Following previous research (15), separate hierarchical linear regressions were conducted to estimate the association between each T1 and T2 inflammatory biomarker and overall hippocampal neurite density. For each model, covariates (maternal education and offspring age, sex, and intracranial volume) were entered in the first step of the model. The second step added the biomarker of interest.

To investigate the effect of PNMI on hippocampal subfield microarchitecture, we repeated our hierarchical linear regressions to estimate the association between each T1 and T2 inflammatory biomarker and neurite density of each hippocampal subregion. For each subregion, four hierarchical multiple regressions were conducted to predict ICVF, one for each biomarker (IL-1RA, IL-6, IL-8, and sTNF-RII, respectively). Since we did not have a priori hypotheses for CA4 and subiculum regions, we applied a multiple comparison correction. To control for multiple comparisons, False Discovery Rate (FDR) correction was applied using the Benjamini-Hochberg (BH) procedure (142). This method controls the expected proportion of false positives while maintaining statistical power (143). P-values were adjusted using the FDR method in R, ensuring appropriate correction for multiple hypothesis testing.

All model results were examined for evidence of and met regression assumptions, including linearity, homoscedasticity, independence between observations, and normality of residuals.

## Acknowledgments

The authors thank the Child Health and Development Studies and Healthy Brains Research Teams and the participating families for their ongoing dedication to our studies. This work was supported by grants NIMH R01 MH118545 and NIMH R01 MH096478 (PI, Ellman), and grants N01HD13334 and N01HD63258 from (PI, Cohn).

