## Supplemental Tables S1 and S2 for "Decreased hippocampal neurite density in late middle-aged adults following prenatal exposure to higher levels of maternal inflammation"

**Table S1.** *Spearman’s correlations for variables of interest: first trimester.*
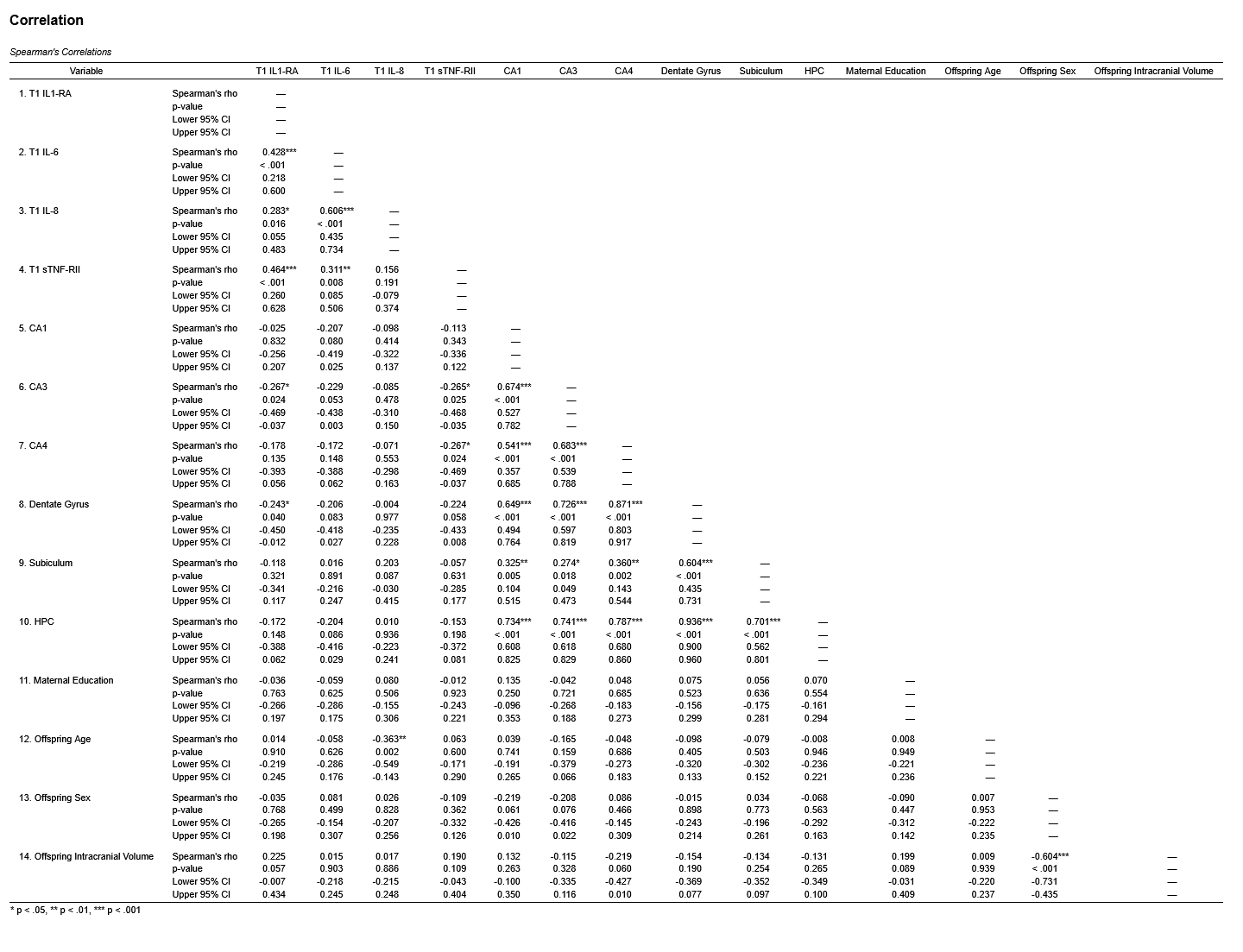

*Note.* Covariates included were maternal education at birth, offspring age, sex, and intracranial volume; IL = Interleukin, RA = receptor antagonist, sTNF-RII = soluble tumor necrosis factor receptor-II, HPC = hippocampus, CA = Cornu Ammonis, Dentate Gyrus includes the molecular and granule cell layers; * *p* < 0.05, ** *p* < 0.01, *** *p* < 0.001

**Table S2.** *Spearman’s correlations for variables of interest: second trimester.*
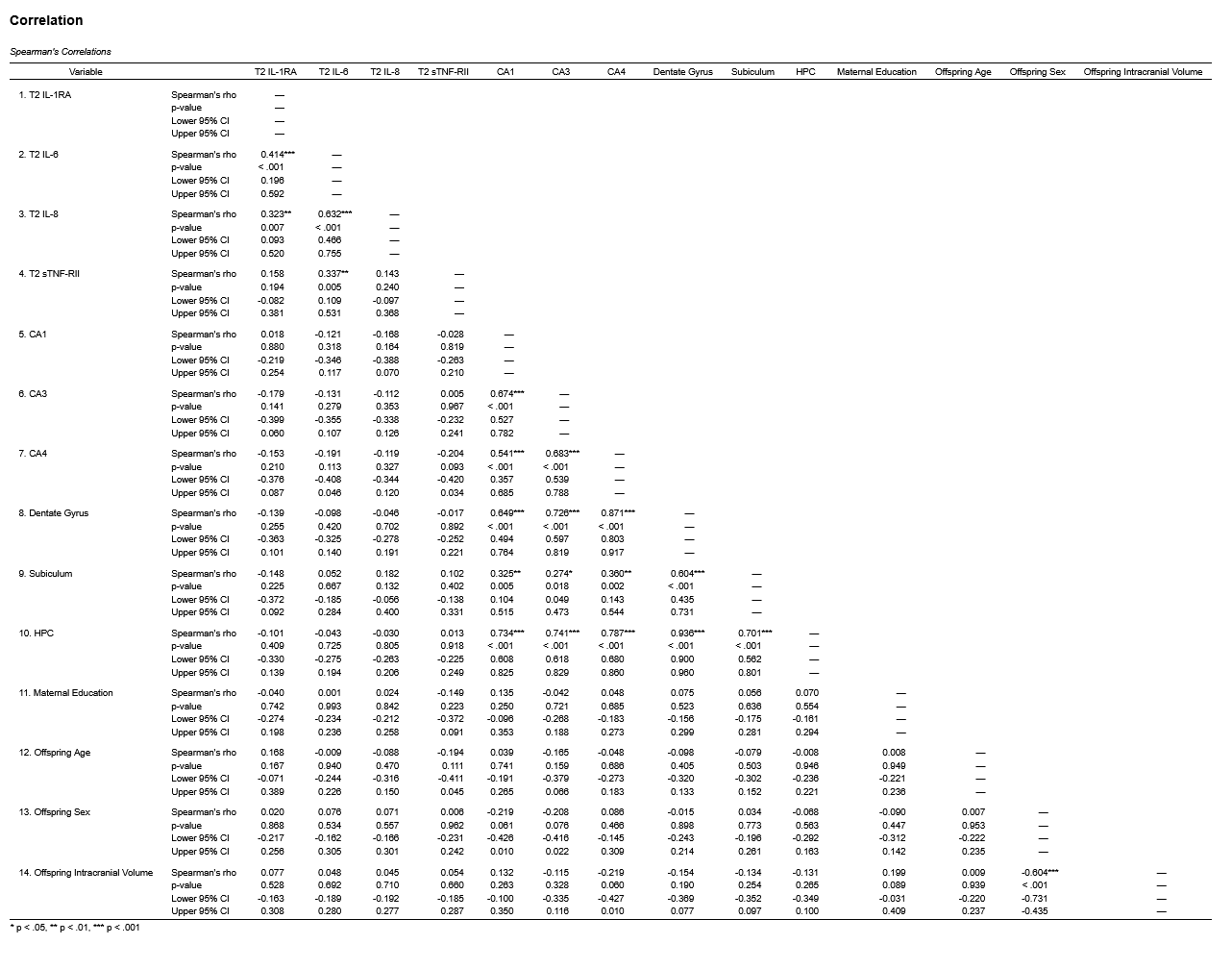

*Note.* Covariates included were maternal education at birth, offspring age, sex, and intracranial volume; IL = Interleukin, RA = receptor antagonist, sTNF-RII = soluble tumor necrosis factor receptor-II, HPC = hippocampus, CA = Cornu Ammonis;Dentate Gyrus includes the molecular and granule cell layers; * *p* < 0.05, ** *p* < 0.01, *** *p* < 0.001
